# Substrate Temperature and Seed Scarification on Germination Parameters of Butterfly Pea (*Clitoria ternatea*)

**DOI:** 10.1101/2022.02.17.480912

**Authors:** Sean M. Campbell, Brian J. Pearson, S. Christopher Marble

**Affiliations:** Department of Environmental Horticulture, University of Florida, Institute of Food and Agricultural Sciences, Mid-Florida Research and Education Center, Apopka, Florida, United States of America

## Abstract

Recent interest in the cultivation and use of butterfly pea (*Clitoria ternatea*) has increased demand for its commercial production. Successful germination of seed is a critical first step to its production; however, few studies have documented factors influential to germination success. To help support commercial production efforts, the influence of substrate temperature and seed scarification technique on germination parameters were evaluated. Two scarification (nicked and nicked and soaked) and three substrate temperature treatments (21, 24, and 27 °C) were imposed on *C. ternatea* seed. Following treatments, six germination parameters were recorded. Data showed seeds maintained at a substrate temperature of 21 °C reached maximal germination (54.8%) at the longest mean germination time (2.61 d) and slowest mean germination rate (0.39 d^-1^). Mechanical scarification by nicking the seed coat with a razor blade prior to soaking the seed for 24 hours reached maximal germination (80.95%) at the lowest mean germination time (2.06 d) and fastest mean germination rate (0.51 d^-1^). Optimal germination was attained when seed was scarified and soaked and then germinated in rockwool substrate maintained at a temperature (T_o_) of 21 °C. Results provide commercial growers with production technique information helpful to fast and efficient germination of *C. ternatea* seed.

## Introduction

Butterfly pea (*C. ternatea*) has long been valued as a forage legume in tropical Asia, China, Sudan, South and Central America and the East and West Indies [1]. More recently, *C. ternatea* has been recognized in the United States and Europe for its ornamental value due to its aesthetically appealing deep blue flowers. Driven by increased interest in the utilization of the flower as a food colorant, nutraceutical, cosmetic, and environmentally friendly insecticide, propagation and production of *C. ternatea* in novel growing environments has been more prevalent than ever [2]. The herbaceous perennial plant generally produces an abundance of seeds (Fig 1). Morris [3] observed a range of between 6 and 3400 total seeds per plant in their assay of 19 different accessions. Despite generally producing a large number of seeds per plant, *C. ternatea* is sometimes regarded for its low seed germination rates.

**Fig 1.**
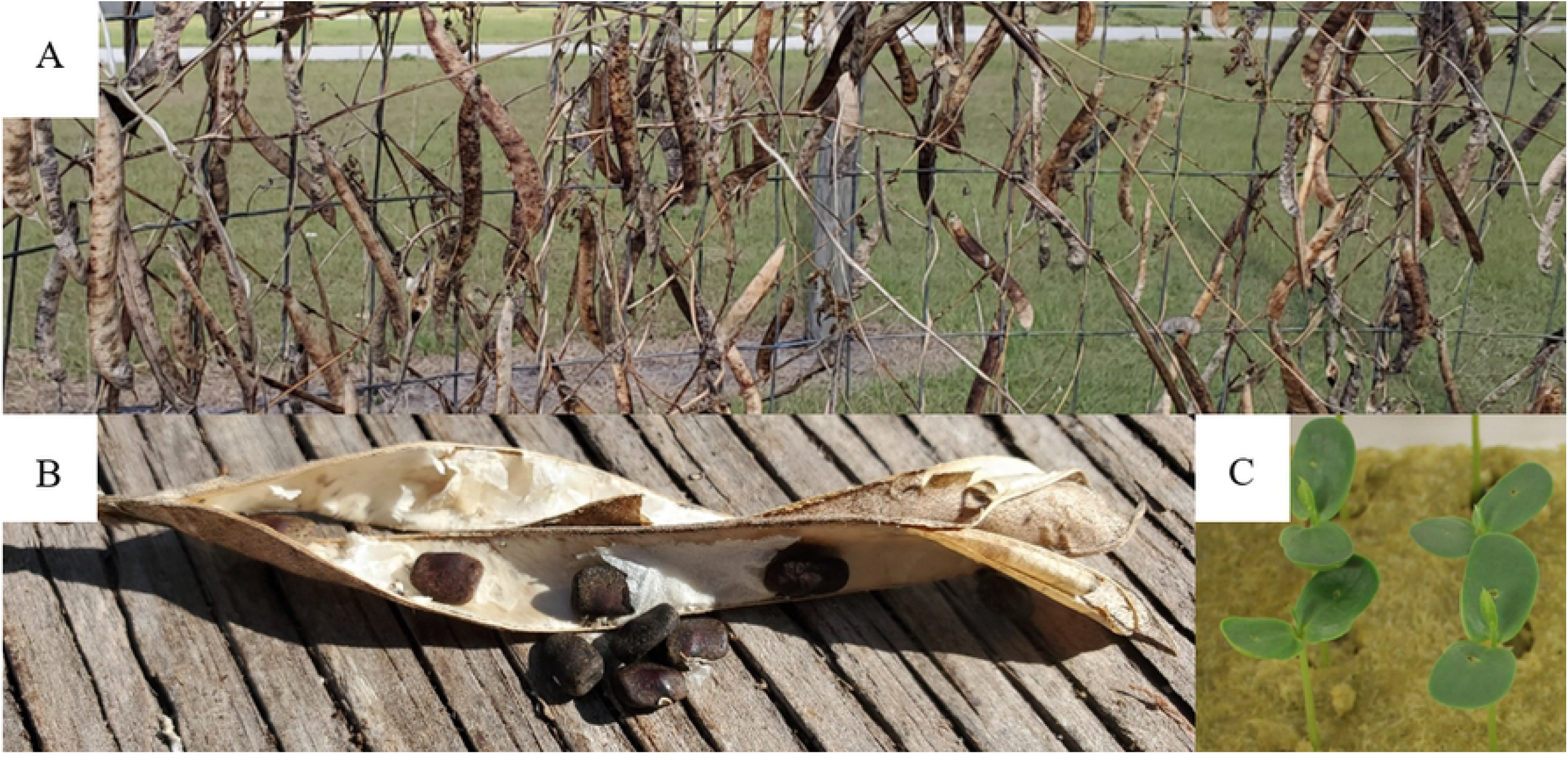
Butterfly pea (*Clitoria ternatea*) seeds and pods. (A) Dried *C. ternatea* seed pods remaining on the vine from the previous year’s growth. (B) *C. ternatea* seeds after being removed from the dried seed pods described. (C) Seven-day-old *C. ternatea* seedlings in soilless substrate under a vented humidity dome.

Seed germination viability and percentage are influenced by a variety of internal and external factors that can have significant implications for commercial production operations. Recent research that examined the influence of substrate type on germination success of *C. ternatea* found rockwool (Grodan, Inc., Roermond, Netherlands) exhibited a higher germination capacity than soilless substrate at 92 and 84% respectively [4]. Substrate temperature is an important external factor given its substantial impact on germination capacity and rate. Varying optimal germination temperatures (T_o_) have been described for *C. ternatea* both as a single temperature of 30 °C [5] and as a range of temperatures of 24 - 28 °C [6] and 24 - 32 °C [2]. Germination capacity (G), also referred to as germinability and described [7] is characterized as the binary response of whether a fully formed cotyledon is visible above the substrate surface at the end of observations (germinated/non-germinated). Recent research examining the influence of substrate temperatures of 21, 24, and 27 °C and G values of 93, 88, and 82% respectively, determined a T_o_ of 21 - 27 °C for *C. ternatea* [4].

Similar to many legumes, *C. ternatea* has a waxy seed coat which is impermeable to water and may result in stunted and inconsistent germination. Freshly harvested *C. ternatea* seeds do not readily absorb water due to their waxy coat [8]. To overcome this, authors stored seed for six months to allow sufficient time for the waxy coat to breakdown naturally which resulted in an increase in germination capacity by 15 - 20%. Despite this increase, the duration of time necessary for increased germination capacity may be in conflict with commercial cultivation objectives and resources where storage space and time limit the ability of aging seed. Several methods exist for increasing the permeability of seed coats with most focused on physical (scarification) or chemical treatments. Mechanical scarification of *C. ternatea* seed through removal of the seed tip has been found to result in a 30% increase in germination capacity (G) over the control [9]. While treatment with sulfuric acid has been shown to be effective for *C. ternatea*, it can result in seedling damage and ultimately cause yellowing of cotyledons following germination [10][1]. To address the need for timely and efficient germination, the influence of three substrate temperatures and two seed scarification methods on germination metrics of *C. ternatea* seed were measured and recorded. Results from this investigation will provide plant producers with critical information necessary to successfully germinate *C. ternatea*.

## Methods

### Seed scarification

Two experiments were performed utilizing the same experimental design. The first experiment occurred 14 – 19 March 2019 and the second occurred 17 – 22 April 2019. For each experiment, 252 seeds were harvested from a single plant grown in a 31 × 15 m gutter-connected greenhouse with 30% light reducing polycarbonate paneling, located in Apopka, FL (lat. 28.64°N, long. 81.55°W). On 14 March and 17 April seeds for each experiment were treated with one of two seed scarification treatments described by [11]; either physical scarification with a small nick to the seed coat using a razor blade (Scar.) or physical scarification with the razor blade before a 24-hour soak in deionized water (DI) water (S&S). To evaluate the effect of scarification technique on germination, a control treatment was incorporated where seeds were neither scarified nor soaked.

### Substrate

Rockwool was utilized for both experiments given its documented ability to support the successful germination of *C. ternatea* [4]. On 15 March and 18 April, eighteen 14-count rockwool cube sheets were prepared by cutting 98-count cube (25 cm^3^) sheets into six pieces, discarding the excess. Cubes were soaked for 30 minutes in water adjusted to pH 5.5 using a pH down buffering solution (General Hydroponics Inc., Sebastopol, CA, United States) before a single *C. ternatea* seed was sown into each plug ∼0.6 cm below the substrate surface. The eighteen substrate sheets were then placed into a propagation unit, six per substrate temperature treatment, as described below.

### Substrate temperature

The propagation unit utilized in this experiment consists of a multi-tier shelfed structure (Compact SunLite 3-Tier Garden; Gardener’s Supply Company, Burlington, VT, United States), with each of the three shelves equipped with a 19 cm vented humidity dome (HydroFarm, Petaluma, CA, United States), 25 × 53 cm heat mat (Vivosun, ShangHai, China), fluorescent lighting (T5; Sunblaster, Langley, BC, Canada), and a mini thermo-hygrometer (Mondi, Vancouver, BC, Canada) to measure environmental conditions within each humidity dome. A digital hygrometer and temperature monitor (AcuRite, Lake Geneva, WI, United States) was placed on the exterior of the propagation unit to record ambient environmental conditions, with mean temperature and humidity of 22.4 °C (53%) and 22.2 °C (50%) for experiments 1 and 2, respectively. Heating mats were programmed to maintain 21, 24, and 27 °C for the three experimental substrate temperature treatments imposed in the study. Top and side vents of the domes were fully closed to maintain consistent humidity. Lights were positioned ∼5 cm from the top of the dome and were run continuously throughout the duration of the experiment, providing photosynthetically active radiation (PAR) at an intensity of between 55 and 90 μmol·m^-2^·s^-1^ at the outer corners and the center of the substrate sheet, respectively. Each of the substrate sheets had approximately 50 mL of water applied each day utilizing a laboratory wash bottle (Thermo Fisher Scientific Inc., Waltham, MA).

### Germination parameters

Six germination parameters were recorded and calculated throughout the experiment [7]. The qualitative value of germination capacity (G), as described above, was then converted into a quantitative attribute of percentage (%) germinability for statistical analysis and reporting. Mean germination time (MT) is the mean of the individual germination times, weighted by the number of seeds germinated per data measurement time interval (days). Mean germination time provides a quantifiable assessment of the average amount of time necessary for maximum germination of an experimental group. Use of the weighted mean for this measurement accounts for variance in the number of seeds germinated per time interval. The coefficient of variation of germination time (CV_t_) is a measurement of the variability in relation to the MT, allowing for additional comparisons independent of the mean germination time magnitude.

Calculated as the reciprocal of MT, the mean germination rate (MR) quantifies germination rate increases and decreases in relation to 1/MT rather than MT alone. Uncertainty of the germination process (U) is a measurement of the degree of uncertainty associated with the frequency and distribution of germination within an experimental group. Since only one seed germinating can change U, this measurement value quantifies the degree of spread of germination as influenced by temporal factors. Conversely, the synchrony of the germination process (Z) represents the degree of overlap that exists during germination and is only produced when two or more seeds finish germination within the same time interval [7].

### Experimental design

The experiment was organized as a 2 × 3 factorial in a completely randomized design, with substrate temperature and seed scarification assigned as independent variables. Four replicates were cultivated for each treatment combination, two per experimental trial, for a total of 36 experimental units. Heating mat temperature (°C), vented humidity dome interior temperature (°C) and humidity (%), and ambient laboratory temperature (°C) and humidity (%) were recorded daily for the portions of the trial where active germination was taking place; a total of six measurements each for Trial 1 (14 - 19 March) and Trial 2 (17 - 22 April) (Table 1). Germination parameters were analyzed based on a collected response then reported as percentages. A restricted maximum likelihood mixed model analysis was performed on the data collected from the 14 germinated seeds per experimental trial using JMP® Pro 14 (SAS; Cary, NC) with post-hoc mean separation tests performed using Tukey’s honest significant difference test by germination trial with variance within treatment combination replicates defined as the random error term. Statistical tests were considered significant if *P* ≤ 0.05.

**Table 1.** Recorded substrate temperature (°C), vented humidity dome interior temperature (°C) and humidity (%) for experimental trials 1 and 2 of the 21, 24 and 27 °C ^z^ substrate temperature treatments.

| Temperature (°C)<br>Treatment | Recorded Substrate<br>Temperature (°C) |  | Interior<br>Temperature (°C) |  | Interior Humidity (%) |  |
| --- | --- | --- | --- | --- | --- | --- |
|  | Trial 1 | Trial 2 | Trial 1 | Trial 2 | Trial 1 | Trial 2 |
| 21 | 24.2<sup>z</sup> | 24.8 | 23.6 | 25.3 | 62.8 | 54.7 |
| 24 | 24.4 | 25 | 23.6 | 25.3 | 62.2 | 55.3 |
| 27 | 26.8 | 26.2 | 75 | 25.6 | 63.3 | 59.3 |

<sup>z</sup> Mean of the six environmental parameters collected daily for Trial 1 (14 - 19 March) and Trial 2 (17 - 22 April).

## Results and Discussion

An interaction effect between substrate temperature and seed scarification was not observed, thus independent variables were analyzed separately (Table 2). G values for the substrate temperature variable lacked difference, but an inverse relationship was observed between increasing substrate temperature and associated G values, with 54.8, 50.0, and 48.8% for the 21, 24, and 27 °C treatments, respectively. Conversely, the scarified and soaked treatment exhibited a higher germination capacity (80.9%) than the scarified (45.8%) and control (26.8%) treatments (*P* < 0.01) (Figs 2 and 3). This is supported by values for the interaction effect; despite lacking significance, the 21 °C S&S, 24 °C S&S, and 27 °C S&S treatments had maximal germination at 82.1%, 82.1%, and 78.6%, respectively (Fig 4).

**Table 2.** Mean germination capacity (G), mean germination time (MT), coefficient of variation of the germination time (CV_t_), mean germination rate (MR), uncertainty of the germination process (U), and synchrony of the germination process (Z) for the 21, 24 and 27 °C substrate temperature independent variables; the scarified, scarified and soaked, and control seed scarification independent variables; and the substrate temperature (°C) by seed scarification interaction effect for butterfly pea (*Clitoria ternatea*) seeds.

| Temperature × Scarification | G (%) |  | MT (d) |  | CV<sub>t</sub> (%) |  | MR (d<sup>-1</sup>) |  | U |  | Z |  |
| --- | --- | --- | --- | --- | --- | --- | --- | --- | --- | --- | --- | --- |
|  | *P* = 0.71 |  | *P* = 0.79 |  | *P* = 0.12 |  | *P* = 0.77 |  | *P* = 0.81 |  | *P* = 0.75 |  |
| 21 °C Control | 26.8 | a | 2.74 | a | 32.2 | a | 0.37 | a | 1.05 | a | 0.21 | a |
| 21 °C Scar.<sup>z</sup> | 55.4 | a | 2.81 | a | 24.0 | a | 0.36 | a | 1.17 | a | 0.41 | a |
| 21 °C S&S | 82.1 | a | 2.29 | a | 22.6 | a | 0.44 | a | 0.99 | a | 0.55 | a |
| 24 °C Control | 25.0 | a | 2.68 | a | 26.3 | a | 0.39 | a | 0.91 | a | 0.32 | a |
| 24 °C Scar. | 42.9 | a | 2.34 | a | 31.5 | a | 0.43 | a | 1.22 | a | 0.36 | a |
| 24 °C S&S | 82.1 | a | 2.20 | a | 40.8 | a | 0.49 | a | 1.40 | a | 0.35 | a |
| 27 °C Control | 28.6 | a | 2.13 | a | 29.4 | a | 0.50 | a | 1.02 | a | 0.34 | a |
| 27 °C Scar. | 39.3 | a | 2.29 | a | 28.4 | a | 0.44 | a | 1.17 | a | 0.44 | a |
| 27 °C S&S | 78.6 | a | 1.71 | a | 49.6 | a | 0.60 | a | 1.12 | a | 0.50 | a |
| Temperature | *P* = 0.53 |  | *P* = 0.02* |  | *P* = 0.14 |  | *P* = 0.01* |  | *P* = 0.84 |  | *P* = 0.67 |  |
| 21 °C | 54.8 | a | 2.61 | b | 26.3 | a | 0.39 | b | 1.07 | a | 0.66 | a |
| 24 °C | 50.0 | a | 2.41 | ab | 32.9 | a | 0.44 | b | 1.18 | a | 0.60 | a |
| 27 °C | 48.8 | a | 2.04 | a | 35.8 | a | 0.52 | a | 1.10 | a | 0.51 | a |
| Scarification | *P* < 0.01* |  | *P* = 0.04* |  | *P* = 0.12 |  | *P* = 0.02* |  | *P* = 0.54 |  | *P* = 0.18 |  |
| Control | 26.8 | c | 2.52 | b | 29.3 | a | 0.42 | b | 0.99 | a | 0.29 | a |
| Scar. | 45.8 | b | 2.48 | a | 27.9 | a | 0.41 | b | 1.19 | a | 0.40 | a |
| S&S | 80.9 | a | 2.06 | a | 37.7 | a | 0.51 | a | 1.17 | a | 0.47 | a |

<sup>z</sup> Scar. = Scarified; S&S = Scarified and soaked
\* = Significance at $P \leq 0.05$ ; means (n = 36 per experimental trial) within column with the same letter are not significantly different ( $P \leq 0.05$ ; Tukey's honest significant difference test).

**Fig 2.**
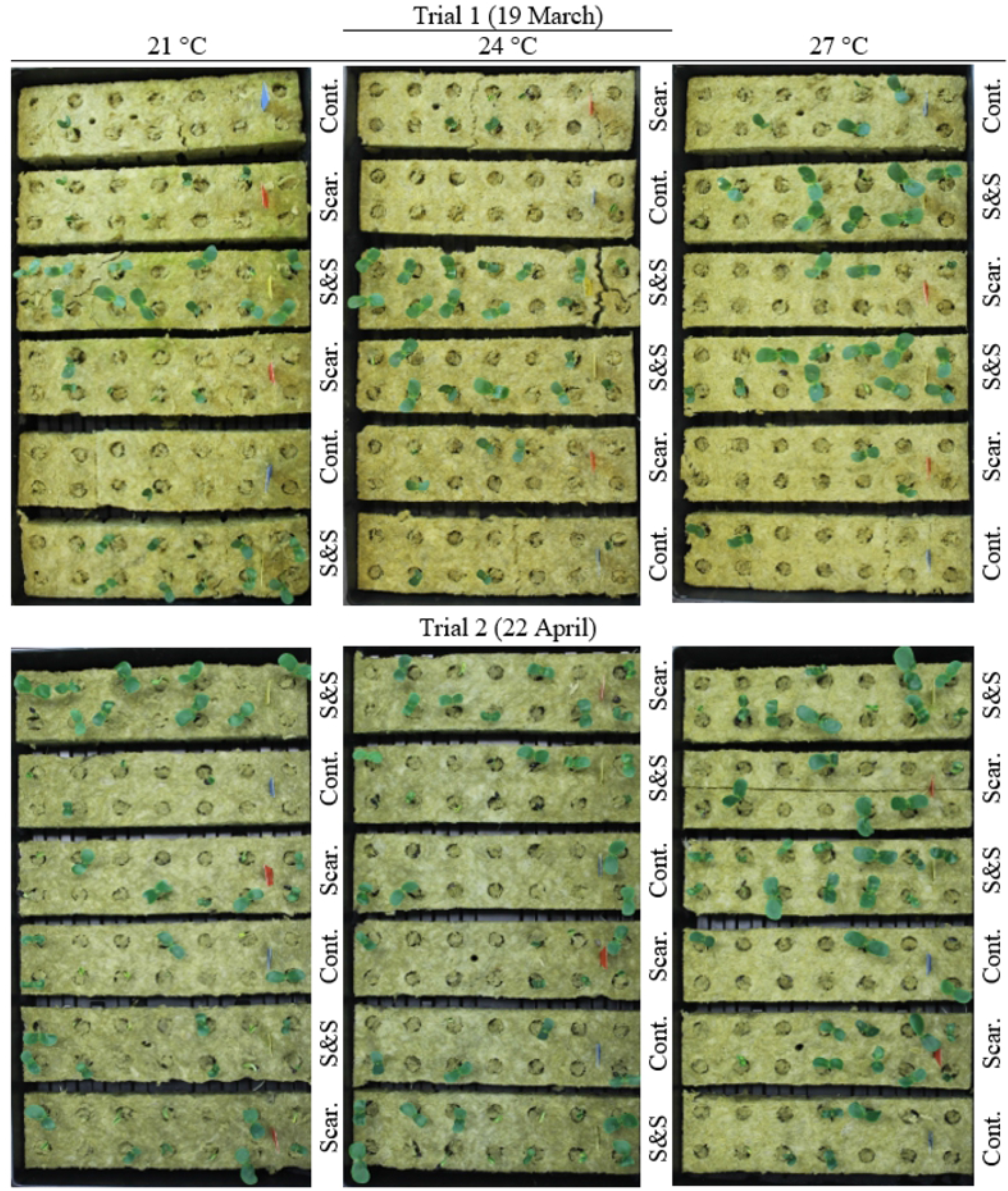
*Clitoria ternatea* experimental design model. Overhead view of germinated butterfly pea (*Clitoria ternatea*) seeds inside the replicate 14-count rockwool plug sheets on the final days of experimental trials 1 and 2. Contains the scarified (Scar.), scarified and soaked (S&S) and control (Cont.) seed scarification treatments nested within the 21, 24, and 27 °C substrate temperature treatments.

**Fig 3.**
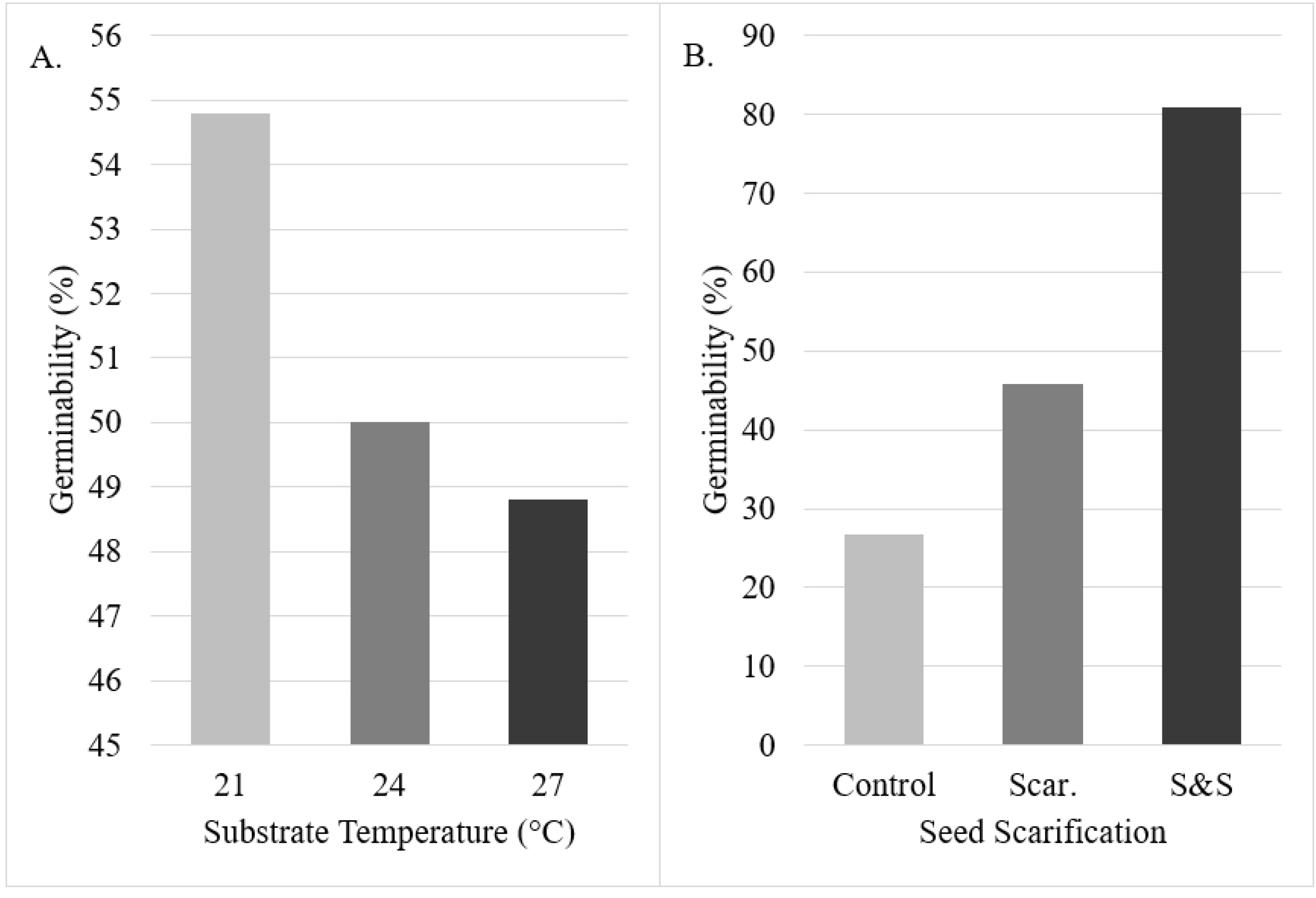
Germinability (%) of butterfly pea (*Clitoria ternatea*) seeds. (A) The influence of 21, 24, and 27 °C substrate temperature on germinability. (B) The influence of control, scarified (Scar.), and scarified and soaked (S&S) seed scarification treatment on germinability.

**Fig 4.**
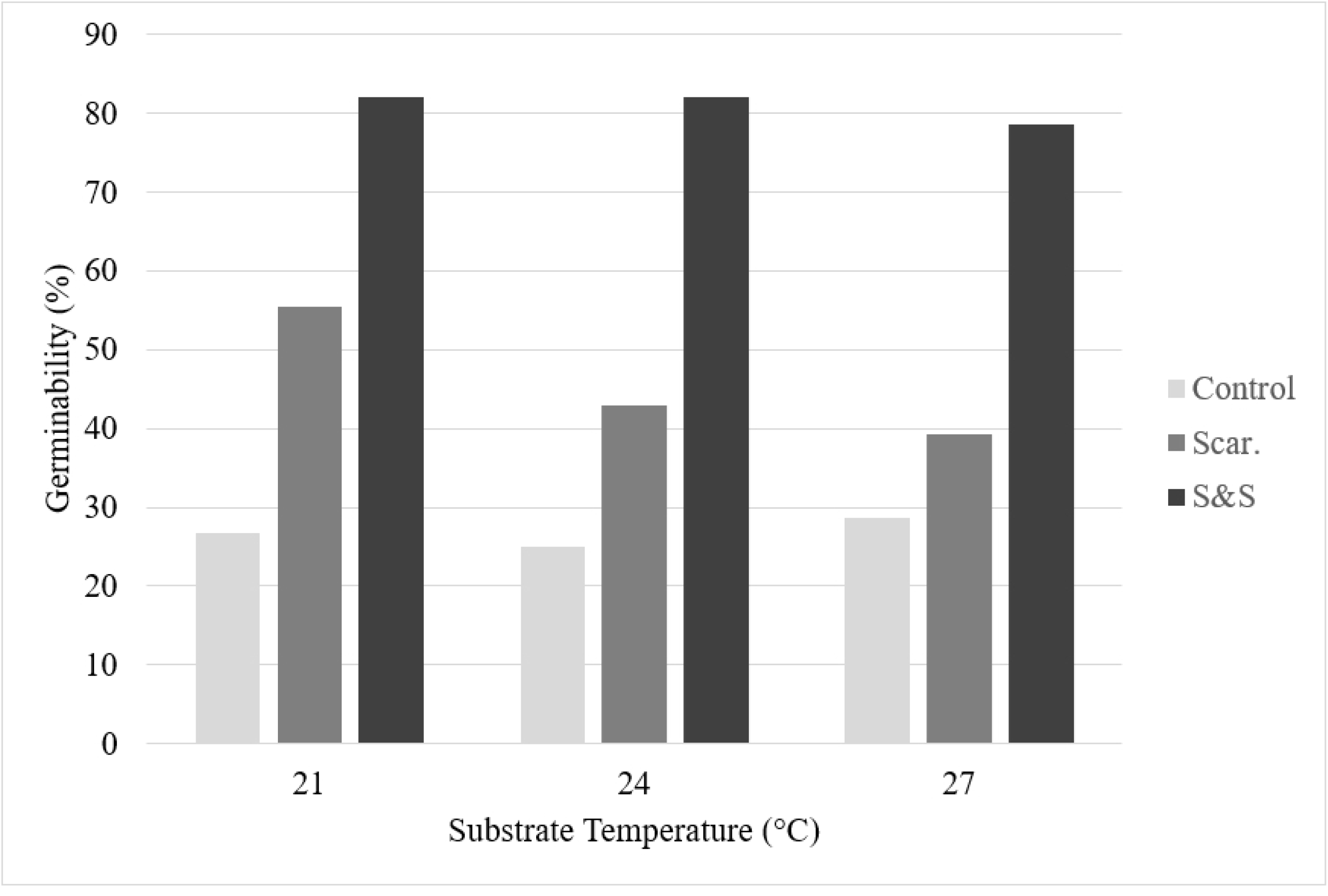
Interaction effect germinability (%) of butterfly pea (*Clitoria ternatea*) seeds. Interaction effect between the 21, 24, and 27 °C temperature treatments and the control, scarified, and scarified and soaked seed scarification treatments.

Differences were also observed for MT and MR among both temperature and scarification treatments (Table 2). MT decreased with increasing substrate temperature for the 21 (2.61 d), 24 (2.41 d), and 27 °C (2.04 d) treatments (*P* = 0.02). MR values rose with increasing substrate temperatures for the 21 (0.39 d^-1^), 24 (0.44 d^-1^), and 27 °C (0.52 d^-1^) treatments (*P* = 0.01). Among scarification treatments, the scarified and soaked (2.06 d) and scarified (2.48 d) groups had lower MT values than the control (2.52 d) (*P* = 0.04). Similarly, the scarified and soaked group (0.51 d^-1^) had a faster germination rate than seed subjected to the scarified (0.41 d^- 1^) or control (0.42 d^-1^) treatments (*P* = 0.02). The interaction effects support these results; with the exception of the 27 °C control group outlier, the 27 °C S&S (1.71 d, 0.60 d^-1^), 27 °C S&S (2.20 d, 0.49 d^-1^), and 21 °C S&S (2.29 d, 0.44 d^-1^) treatments had the shortest mean germination times and the highest mean germination rates.

Considering depressed MT and elevated MR values represent a briefer and more rapid germination, differences observed in the MT and MR parameters of the substrate temperature variable indicate that the 21 °C treatment group outperformed the 24 °C or 27 °C groups. Similarly, while substrate temperature lacked significance for G, the 21 °C treatment produced the highest percentage of viable seedlings. This T_o_ is slightly lower than values published in related literature and is likely explained by variations in how temperature was recorded (i.e., recording substrate temperature as compared to recording air temperature). While other publications measured local atmospheric temperature and inferred its influence on substrate temperature, this investigation used heating mats with integrated thermostats to accurately measure, record, and maintain substrate temperature [5,6]. Thus, application of reported optimal germination temperature by commercial producers should be carefully implemented based on the media reported in literature.

Differences were observed for G, MT, and MR among seed scarification treatments (Table 2). The scarified and soaked experimental treatment produced a higher percentage of viable, germinated seeds than the scarified or control treatment groups. The scarified and soaked treatment resulted in lower MT and higher MR values than the scarified or control groups, indicating that germination was accomplished in a shorter time and at a faster rate. This conclusion is consistent with published literature where the hard seed coat of *C. ternatea* required either a dormancy period or mechanical/chemical scarification to promote germination [2] with previous research finding mechanical scarification in *C. ternatea* resulted in a 30% increase in G values over the control [9].

Considering MT is a quantifiable value relating to the average time for maximum germination, theoretically optimized germination conditions should result in a shorter germination period and minimal MT value. As the reciprocal, optimized germination conditions should therefore also result in faster average germination rate and maximal MR value. But when considered in the context of the germination capacities, a different trend emerges, with a direct relationship between increasing germination capacity and mean germination time and rate for the substrate temperature variable and the inverse for the seed scarification variable. This is further supported by the substrate temperature and seed scarification interaction effect. With the exception of the 27 °C control (28.6%, 2.13 d), the 21 °C S&S (82.1%, 2.29 d), 24 °C S&S (82.1%, 2.20 d) and 24 °C S&S (78.6%, 1.71 d) treatment groups had the highest G and lowest MT values, respectively. Similarly, with the exception of 27 °C control (28.6%, 0.50 d^-1^) group, the 21 °C S&S (82.1%, 0.44 days^-1^), 24 °C S&S (82.1%, 0.49 days^-1^) and 27 °C S&S (78.6%, 0.60 days^-1^) treatment groups had both the highest G and MR.

The final germination parameters CV_t_, U, and Z lacked significance (*P* = 0.14, 0.84, and 0.67, respectively) (Table 2). With an observed increase in CV_t_ with increased substrate temperature, the 21 °C (26.3%) treatment group had less variability in germination than the 24 (32.9%) or 27 °C (35.8%) treatments. Relationships between CV_t_ and seed scarification were less consistent as the scarified (27.9%) treatment group had more variability than the scarified and soaked (37.7%) or control (29.3%) treatments. Less association was observed for the U and Z metrics; the only values consistent with trends observed for the other germination parameters are the Z values of the seed scarification variable, with the scarified and soaked (0.47) group showing higher synchrony than the scarified (0.40) or control (0.29) groups.

## Conclusion

With a surge of interest in a wide variety of applications, the need to maximize germination viability for *C. ternatea* seed has become increasingly crucial to commercial producers. Both variables of substrate temperature and seed scarification had experimental treatments that resulted in maximized germination capacities but with opposite influences on the MT and MR parameters. The 21 °C substrate temperature independent variable reached maximal germination (54.8%) at the highest mean germination time (2.61 d) and slowest mean germination rate (0.39 d^-1^), while the scarified and soaked seed scarification treatment reached maximal germination (80.9%) at the lowest mean germination time (2.06 days) and fastest mean germination rate (0.51 d^-1^). Regardless of these conflicting trends, optimal germination was achieved through mechanical scarification by nicking the seed coat with a razor blade prior to soaking the seed for 24 hours, followed by germination of seed in rockwool at a substrate temperature (T_o_) of 21 °C. Research efforts that examine additional environmental factors responsible for germination success, such as those that occur naturally outdoors, is warranted and likely helpful to support commercial cultivation of *C. ternatea*.

## Acknowledgements

We would like to acknowledge Mengzi Zhang and Caroline Warwick for their scientific and technical assistance. Additionally, we would like to recognize Roger Kjelgren for providing administrative support and encourgement to pursue this research.

## References

1. Awadalla, A Morsy, A. Growth, forage yield and quality of clitoria (Clitoria ternatea) as affected by dates and methods of sowing and phosphorus fertilizer under Toshka region condition. Middle East J. of Agr. Res. 2017; 6:506–18.

2. Oguis, G, Gilding, E, Jackson, M, Craik, D. Butterfly pea (Clitoria ternatea), a cyclotide-bearing plant with applications in agriculture and medicine. Frontiers in Plant Sci. 2019; 10. doi:10.3389/fpls.2019.00645

3. Morris, J. Characterization of butterfly pea (Clitoria ternatea L.) accessions for morphology, phenology, reproduction and potential nutraceutical, pharmaceutical trait utilization. Genet. Resources and Crop Evolution. 2009; 56:421–27. doi:10.1007/s10722-008-9376-0

4. Campbell, S, Pearson, B, Marble, S. Substrate type and temperature on germination parameters of butterfly pea. HortTechnology. 2020; 30:398–403. doi:10.21273/HORTTECH04583-20

5. Selvamaleeswaran, P, Wesely, J, Vennila, B, Balakrishnan, S. Dormancy breaking and seed germination techniques for Clitoria ternatea Linn. Intl. J. of Appl. Biotechnol. and Biochem. 2011; 1:185–92.

6. McDonald, C. Germination response to temperature in tropical and subtropical pasture legumes. 1. Constant temperature. Austral. J. of Expert. Agr. 2002; 42:407–19.

7. Ranal, M, Santana, D, Ferreira, W, Mendes-Rodrigues, C. Calculating germination measurements and organizing spreadsheets. Brazilian J. of Bot. 2009; 32:849–55.

8. Mullick, P, Chatterji, U. Eco-physiological studies on seed germination: Germination experiments with the seeds of Clitoria ternatea Linn. Trop. Ecol. 1967;8:117–25.

9. Mishra, D, Chaudhuri, K, Singh, V, Shukla, J. Seed germination studies in medicinal plants of commercial value. In: Tewari, V, Srivastava, R, editors. Multipurpose Trees in the Trop.: Mgt. & Improvement Strategies. Jodhpur: Scientific Publishers; 2006. p. 382.

10. Makasana, J, Pillai, V, Sharma, A, Dholakiya, B, Gajbhiye, N, Saravanan, R. Effect of seed treatment on germination and flavonoids diversity in accessions of butterfly pea (Clitoria ternatea). Indian J. of Agr. Sci. 2016; 86(12):1553–8.

11. Mackay, W, Davis, T, Sankhla, D. Influence of scarification and temperature on seed germination of Lupinus arboreus. Seed Science and Technol. 1996; 23(3):815–21.

